# Intracranial Validation of Magnetoencephalography Across Oscillatory Frequency and Depth

**DOI:** 10.64898/2026.08.18.745459

**Authors:** Chetan Gohil, George O’Neill, Gareth Barnes, Vladimir Litvak, Mark Woolrich, Shikun Zhan, Wei Liu, Bomin Sun, Chunyan Cao, Daniel Bush, Umesh Vivekananda

## Abstract

Non-invasive, whole-brain neuroimaging methods such as functional magnetic resonance imaging, electroencephalography (EEG), and magnetoencephalography (MEG) are essential tools for studying the basis of human cognition in health and disease. MEG offers the opportunity to study neural activity at its intrinsic timescale, by recording the magnetic fields generated by electrical currents within the brain from outside the skull. Moreover, recently developed optically pumped magnetometers (OPMs) allow these recordings to take place in new settings, for example during naturalistic behaviour and in previously inaccessible populations. These breakthroughs have led to a shift in the neuroimaging landscape, with a global increase in the adoption of MEG. Crucially, however, the extent to which MEG recordings can measure different features of neural activity remains unclear. To address this issue, we leveraged a unique and rare dataset of concurrent MEG and intracranial EEG recordings from a cohort of epileptic patients. We found that group-level inferences of spontaneous oscillatory dynamics made with source-localised MEG, i.e. estimates of power and bursts, accurately reflected the underlying neural activity. As expected, the agreement was strongest for lower-frequency activity (delta, theta, and alpha) and superficial sources, and weakest in the gamma range. Crucially, however, MEG was also sensitive to deep structures: it captured oscillatory power and burst dynamics in the hippocampus, most robustly in the theta band. These findings demonstrate that MEG is sensitive to physiologically meaningful activity in cortical and subcortical regions and establish a foundation for the interpretation of future MEG studies across a wide range of research domains.

## Introduction

Neuroimaging is central to understanding the mechanistic basis of human cognition in health and disease. The available modalities each balance spatial and temporal resolution: functional magnetic resonance imaging (fMRI)^1^ offers high spatial resolution but measures a slow haemodynamic proxy for neural activity, while scalp electroencephalography (EEG)^2^ recovers fast neuronal dynamics but is attenuated and blurred by the skull and scalp, reducing its sensitivity to low-amplitude sources and limiting its spatial resolution. Magnetoencephalography (MEG)^3^ occupies a valuable position among these methods, recording the magnetic fields generated by neuronal currents from outside the head at the natural, millisecond timescale of neural activity and with whole-brain coverage^4,5^. Because biological tissue is largely transparent to these magnetic fields, MEG is less distorted than EEG by the intervening skull and scalp. Recently developed optically pumped magnetometers (OPMs) further extend these advantages: without the need for cryogenic cooling, these sensors can be placed directly on the scalp in lightweight arrays, improving the signal-to-noise ratio and enabling recordings during naturalistic behaviour^6–9^ and from previously inaccessible populations^10^. Together, these developments have driven a rapid, worldwide adoption of MEG for human neuroscience.

However, realising the full potential of MEG depends on accurately localising the underlying neural sources of the magnetic fields measured from outside the head. This is an inherently ill-posed problem: several configurations of neural sources can give rise to the same magnetic field. A range of source-reconstruction algorithms, most prominently the linearly constrained minimum variance (LCMV) beamformer, are used routinely for this purpose^5,11^, but the fidelity of these reconstructions has yet to be properly validated against direct measurements of the underlying neural activity. The accuracy of MEG source reconstruction is expected to decline with distance from the sensors and with frequency, since deep sources produce weaker external fields and high-frequency activity is lower in amplitude. However, many subcortical structures are central to cognition and disease; the hippocampus, for example, is essential for learning and memory^12,13^ and among the first brain regions to be affected by Alzheimer’s disease^14^. Although simulation studies suggest that deep activity can in principle be recovered with MEG^15^, such predictions lack empirical validation against a ground truth.

Stereoelectroencephalography (SEEG) in clinical populations provides one opportunity to address this issue: surgically implanted electrodes record local field potentials from within the brain with millimetre precision, including from deep structures^16^. However, simultaneous intracranial and MEG recordings from the same brain are rare and technically difficult to acquire. Existing evidence from simultaneous MEG-SEEG is largely confined to individual cases, specific structures, or epileptiform and task-evoked responses rather than spontaneous, ongoing activity^17,18^. Consequently, it remains unclear which features of spontaneous neural activity, such as oscillatory power and transient bursts, can be faithfully recovered by MEG. Here, we answer this question using a rare, large simultaneous MEG-SEEG dataset acquired at rest (*N*=14; 7 female; 18-47 years old). Specifically, we quantify which features of spontaneous neural activity can be reconstructed from MEG recordings, and how this varies as a function of oscillatory frequency and cortical depth.

## Results

### MEG correlates with SEEG, most strongly for superficial regions and low frequencies

Simultaneous recordings provide a direct test of how much information is shared between MEG and SEEG. To quantify this correspondence, we used canonical correlation analysis (CCA) between all MEG sensors and the SEEG contacts in each brain region, separately for six canonical frequency bands, assessing significance with a block-shuffling permutation procedure. CCA finds the maximum correlation between a set of MEG and SEEG recordings, where higher values imply they reflect a common signal. Significant canonical correlations were observed between the MEG and SEEG signals across the cohort (Figure 1D), and the same was true when the analysis was repeated on source-reconstructed virtual electrodes (VEs) placed at the SEEG contact locations estimated using a beamformer (Figure S1). These results confirm that the two modalities have correlated dynamics, consistent with previous reports of correspondence between MEG and intracranial recordings based on smaller cohorts^19^. As expected, the strength of the correlation depended systematically on both the depth of the SEEG contacts and the frequency band. Canonical correlations decreased with contact depth, and this depth dependence was significant for delta, theta, alpha, and beta (delta: *r* = −0.29; theta: *r* = −0.29; alpha: *r* = −0.30; beta: *r* = −0.30; all *p* < 0.05) but not for the gamma range (Figure 1D). Canonical correlation was therefore strongest for superficial regions and low frequencies, although some shared information remained for deeper contacts, particularly in the delta/theta range, in line with the reduced (but non-zero) sensitivity of MEG to deep sources reported previously^20,21^.

**Figure 1:**
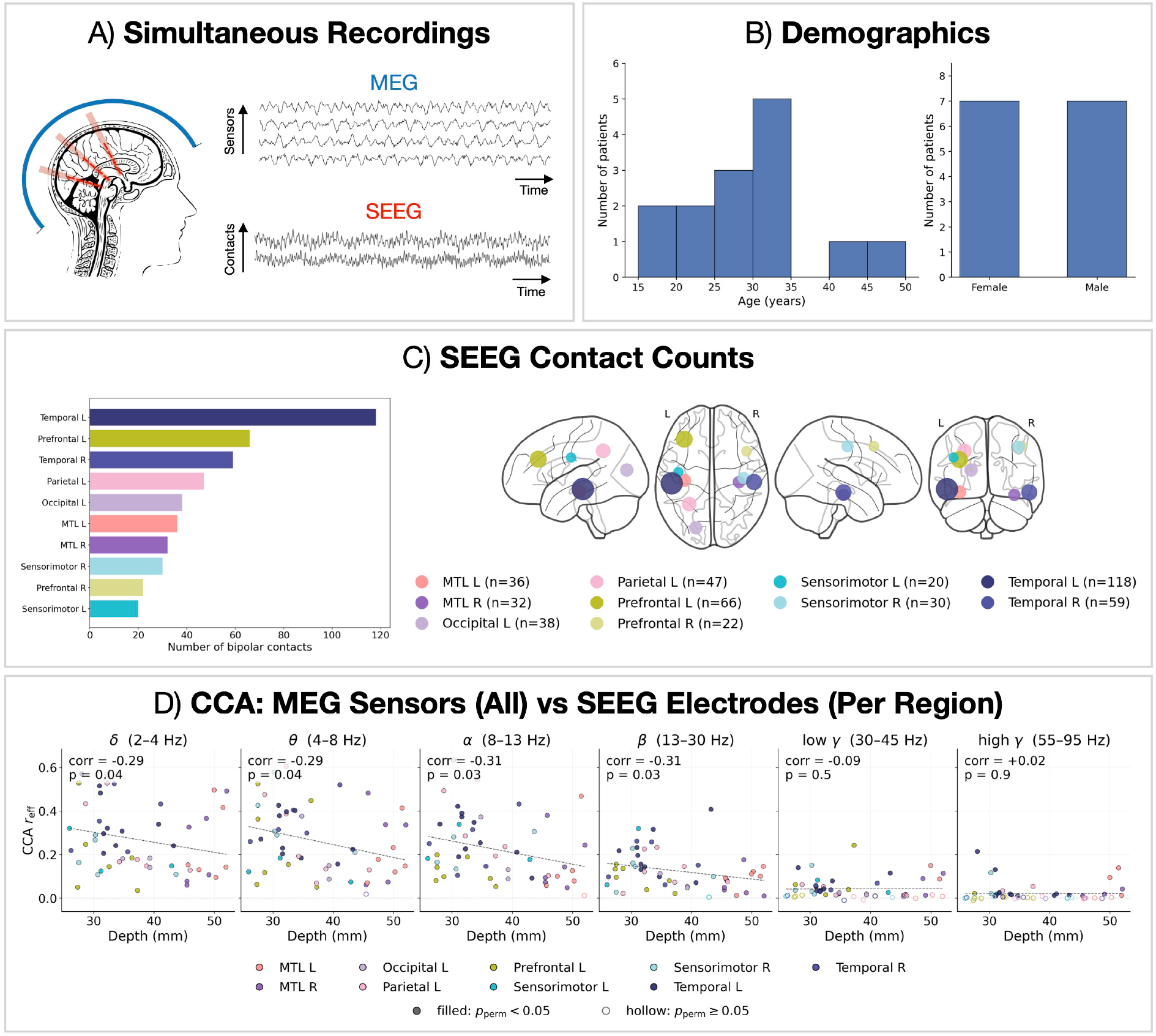
Simultaneous MEG-SEEG dataset. A) Schematic of the simultaneous recording setup: whole-head MEG (sensors) and SEEG (depth-electrode contacts) were acquired concurrently at rest, with illustrative MEG (top) and SEEG (bottom) time series shown. B) Age and sex distribution of the *N* = 14 participants (18-47 years old). C) Number of bipolar SEEG contacts per anatomical region (left) and the corresponding electrode locations on a glass brain (right), coloured by region; only regions that enter the region-level analyses (≥ 3 sessions and ≥ 10 bipolar contacts pooled) are shown. D) Canonical correlation analysis (CCA) between all MEG sensors and the SEEG contacts of each region, plotted against mean contact depth for six frequency bands. Each point is a region in one session, coloured by region; the *y*-axis shows the effective CCA correlation (CCA*r*_eff_). Filled points denote a significant canonical correlation (block-permutation test, *p*_perm_ < 0.05).

### Oscillatory power in MEG correlates with SEEG across sessions

Having established that the band-limited MEG and SEEG signals correlate, we next asked which specific oscillatory features are shared between them. To do so, we computed the power spectral density (PSD) for each region, as well as the band-limited power from each SEEG contact and the corresponding MEG-VE, and correlated those estimates across sessions (Figure 2). At the group level, the MEG-VE PSDs were systematically flatter than the SEEG PSDs, with relatively higher power above 20Hz and reduced power at low frequencies (Figure 2A). This pattern is consistent with the MEG-VE signal being a noisier version of the SEEG: indeed, adding white noise to the SEEG reproduced the same spectral flattening: because the spectra are normalised (z-scored to unit total variance), the broadband floor raises the total variance and lowers the relative power at low frequencies while raising it at high frequencies (Figure S2). Nevertheless, static (time-averaged) oscillatory power was correlated between the two modalities in every frequency band (Figure 2B; delta: *r* = +0.49; theta: *r* = +0.43; alpha: *r* = +0.39; beta: *r* = +0.75; low-gamma: *r* = +0.57; high-gamma: *r* = +0.27), with the correlation strongest in the beta band and weakest at high-gamma. Moreover, this correspondence was spatially specific: the observed correlations exceeded a null distribution obtained by reconstructing the VEs at random locations (Figure S3, all *p*_FDR_ < 0.05).

**Figure 2:**
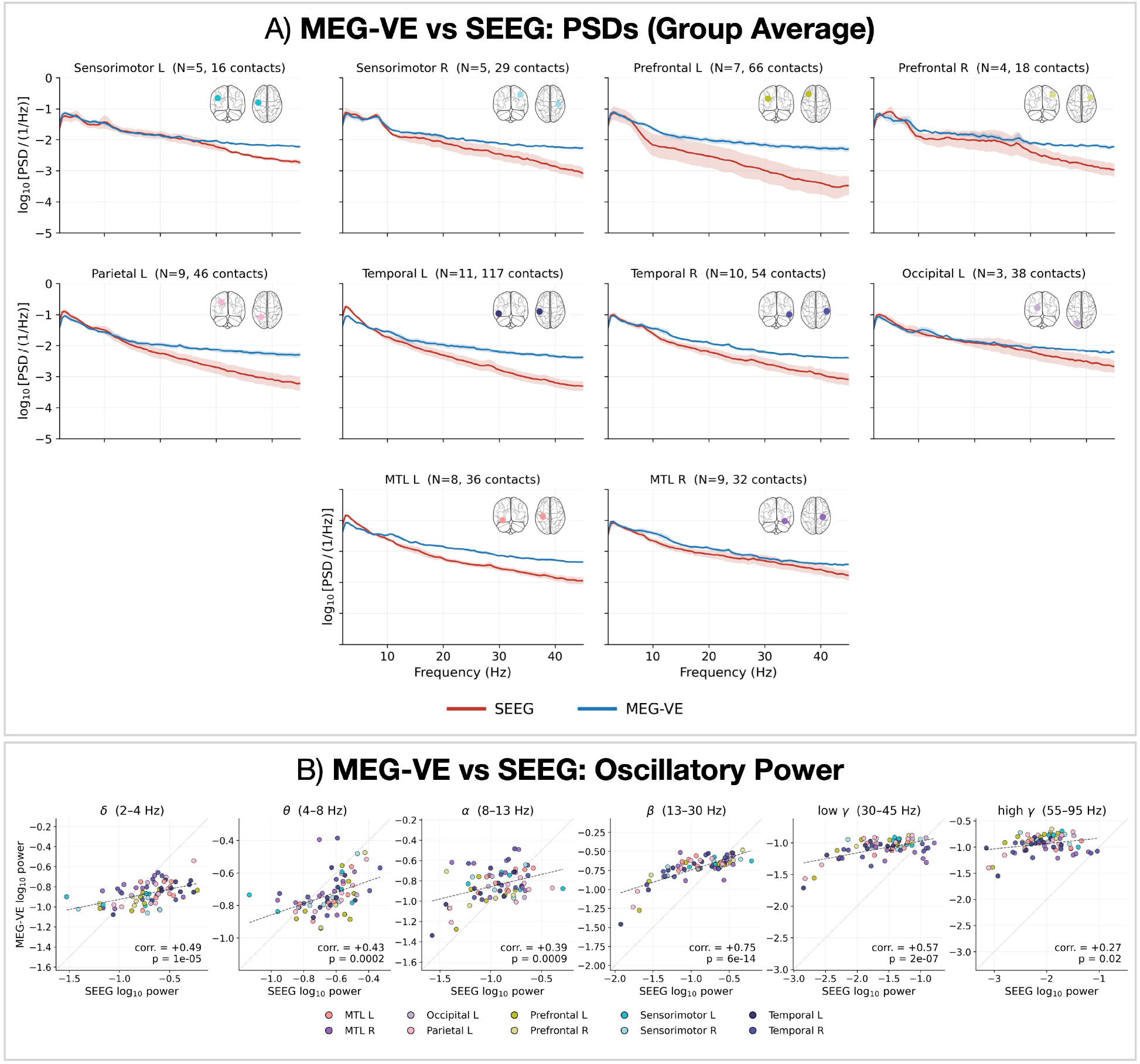
MEG-VE vs SEEG: oscillatory power comparison. A) Power spectral density (PSD; log_10_ scale) computed from MEG virtual electrodes (MEG-VE; blue) and SEEG (red), averaged across sessions and contacts within each region (line, mean; shaded, variability). The number of sessions (*N*) and contacts contributing to each region is annotated and glass-brain insets indicate the region location. B) Across-session correlation between oscillatory power estimated from MEG-VE and SEEG, for six frequency bands. Each point is a region in one session (coloured by region); the dashed line is a straight line fit. Points are pooled across sessions and regions and treated as independent, so the annotated correlation is descriptive; spatial specificity is tested against the random-VE null in Figure S3.

### MEG-detected oscillatory bursts coincide with amplitude increases in SEEG

Next, we asked whether MEG can resolve dynamics in neural activity, in particular, oscillatory bursts, which are transient, high-amplitude events that are widely implicated in cognitive function^22,23^. To test whether such transient dynamics are also captured by MEG, we detected oscillatory bursts in the MEG-VE signals and quantified the accompanying change in SEEG amplitude using an amplitude modulation index (see Methods), computed separately for each frequency band (Figure 3). SEEG amplitude was elevated during MEG-VE bursts of the same band, giving a positive modulation index that was strongest in the theta and alpha bands and approached zero in the gamma range (Figure 3). As with the oscillatory power analysis, this effect was spatially specific, exceeding a random-VE null (Figure S4).

**Figure 3:**
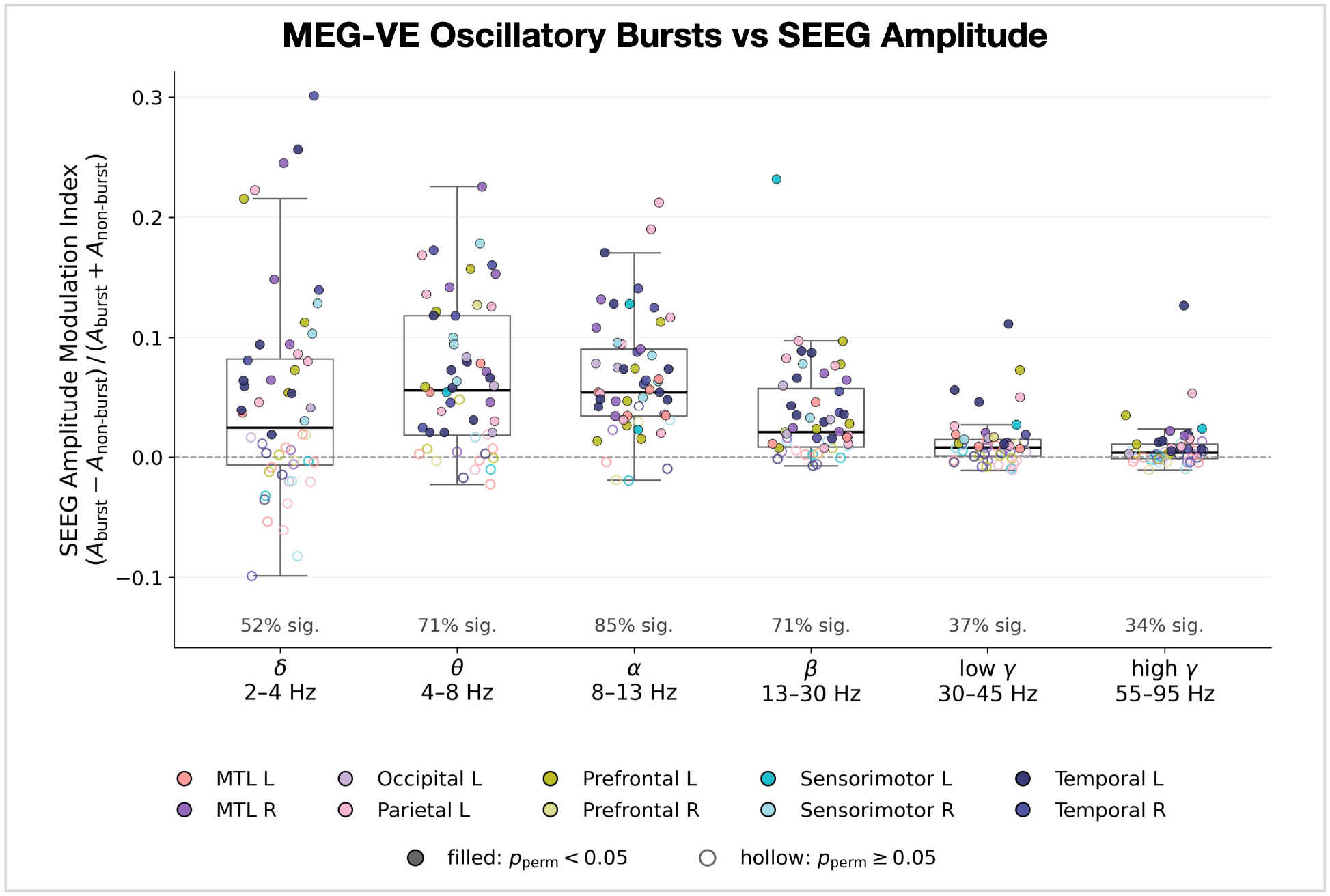
MEG-VE vs SEEG: amplitude modulation in SEEG during oscillatory bursts in MEG. For each frequency band, the SEEG amplitude modulation index quantifies the change in SEEG amplitude during oscillatory bursts detected in the simultaneously recorded MEG-VE. Each point is a region in one session (coloured by region); box plots show the distribution across sessions and filled points denote a significant modulation (block-permutation test, *p*_perm_ < 0.05).

### MEG is sensitive to oscillatory dynamics in the hippocampus

Finally, we asked whether MEG is sensitive to oscillatory activity in a subcortical brain region of particular interest, the hippocampus. This structure is widely assumed to lie beyond the reach of MEG: it sits deep in the brain, and its curved geometry is thought to produce partially cancelling fields that are weak at the sensors^24^. To do so, we examined VEs placed at hippocampal contacts (Figure 4). These MEG-VE captured key features of the SEEG signal, with oscillatory power estimates correlating with the SEEG across sessions in the delta, theta, and beta bands, though not for alpha or the gamma range (Figure 4A). This power correspondence was spatially specific to the hippocampus: for delta, theta, and beta the observed correlation exceeded a depth-matched random-VE null (Figure S5). The MEG-VE also captured the amplitude dynamics of hippocampal bursts. Applying the same amplitude modulation index used across the brain (Figure 3) to hippocampal contacts, the simultaneously recorded SEEG amplitude was elevated during bursts detected on the hippocampal MEG-VE for the theta, alpha, and beta bands, with a trend in delta (Figure 4B). This modulation was spatially specific to the hippocampus in the alpha band, where it exceeded a matched-depth null with a theta trend, whereas the delta modulation was reproduced by the matched-depth null and therefore reflects a spatially broad rather than hippocampus-specific effect (Figure 4B). These hippocampal findings are consistent with prior reports that MEG can capture deep and mesial activity^15,17,25,26^ and with deep-source validation using OPM-MEG and optogenetics^8,27^. Our results suggest that MEG is sensitive to hippocampal activity, particularly in the theta band.

**Figure 4:**
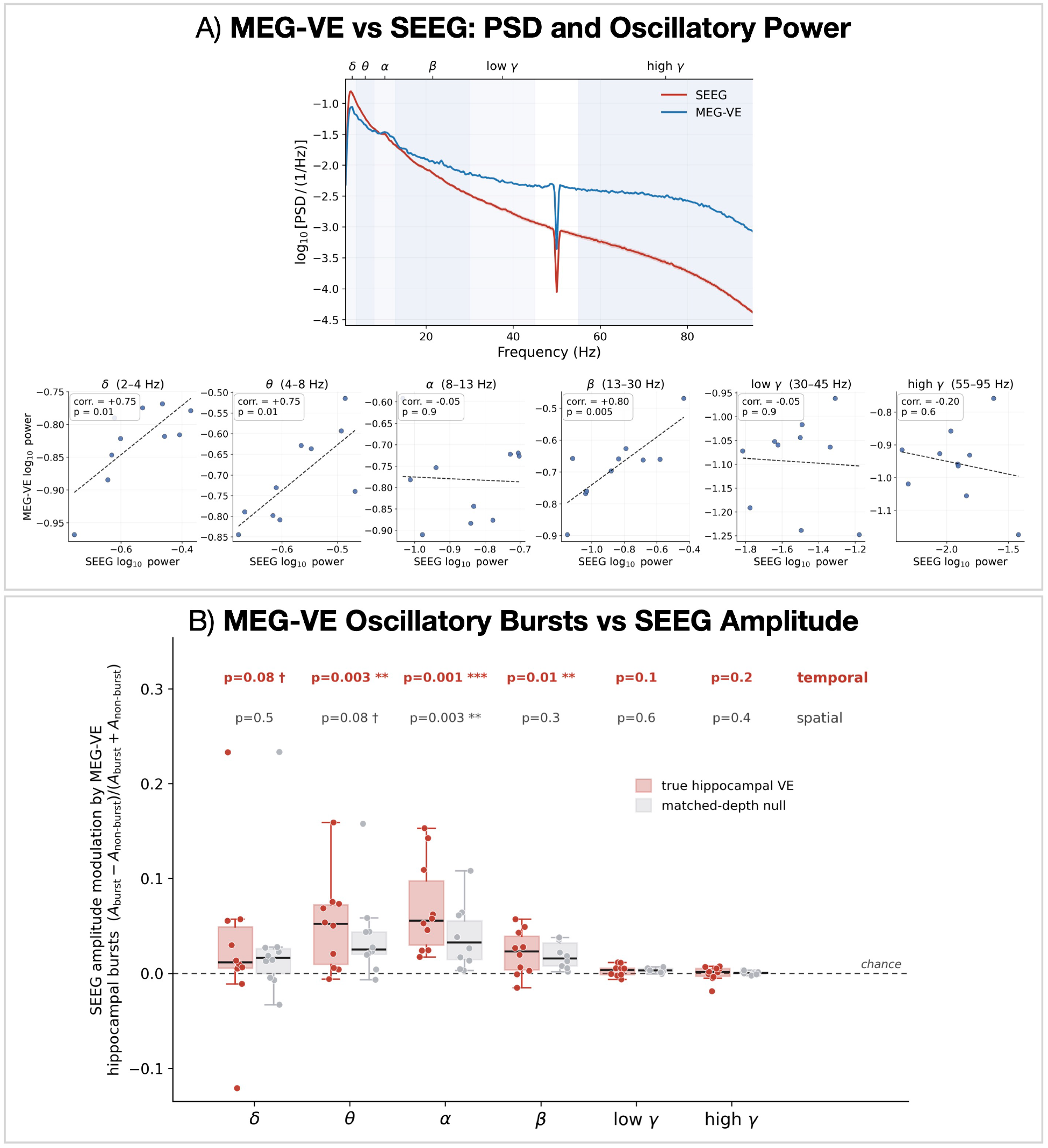
Hippocampal source reconstruction with MEG. A) Power spectral density (top; SEEG red, MEG-VE blue, with frequency bands shaded) and across-session correlation of oscillatory power (bottom) for hippocampal contacts. B) Amplitude modulation of the SEEG during hippocampal MEG-VE bursts. For each band, the SEEG amplitude modulation index for bursts detected on the hippocampal MEG-VE (red) and on matched-depth null VEs (grey); each point is a session. Values above the dashed chance line (zero) indicate temporal coupling (SEEG amplitude elevated during MEG-VE bursts); the true (red) box above the matched-depth (grey) box indicates spatial specificity to the hippocampus. Above each band, the temporal (modulation vs chance) and spatial (true vs matched-depth null) one-sided Wilcoxon *p*-values are annotated.

## Discussion

In this study, we used a relatively large (*N*=14) and rare simultaneous resting-state MEG-SEEG dataset to examine how well source-reconstructed MEG recovers oscillatory activity across the brain and across frequencies. We found that MEG does faithfully capture oscillatory power and activity from transient bursts in both cortical and subcortical regions, including the hippocampus. This work extends prior simultaneous MEG-intracranial studies^17,25,28^ that focused on interictal spikes or task-triggered detection of activity to the imaging of ongoing spontaneous activity in a larger cohort. Importantly, source localising spontaneous activity is a more demanding test than the epileptiform discharges and task-evoked responses targeted previously, as it is not time-locked, and so cannot be isolated by averaging or spike-triggering^17,18^.

### Sensitivity of MEG to deep activity

MEG is often believed to have limited sensitivity to deep brain structures^24^; our results nuance this view, showing that sensitivity is reduced with depth but is non-zero and physiologically meaningful, particularly at low frequencies (Figure 1D). A central finding is that the hippocampal oscillatory dynamics captured by the MEG-VE reflect true neural activity, as measured by the simultaneously recorded SEEG (Figure 4). This is consistent with prior reports of detectable deep and mesial MEG signals that focused on time-locked activity using alternative methods for isolating sources: Pizzo et al.^17^ recovered a small but reliable hippocampal and amygdalar contribution to the surface MEG from spike-triggered recordings, and Dalal et al.^25^ captured task-evoked hippocampal theta, with further evidence from beamformed comparisons against depth recordings^26^ and from a memory-related mesial-temporal component^18^. To this literature we add spontaneous (resting-state) oscillatory dynamics.

Because these estimates were obtained by beamforming to each SEEG contact, a key concern is that they could reflect spatial leakage from stronger cortical sources rather than genuine deep activity^29,30^. Several observations indicate otherwise: the cross-modal correspondence exceeded a random-VE null, in which the VEs were reconstructed at spatially displaced locations, for both oscillatory power (Figure S3) and amplitude modulation (Figure S4), and the canonical correlation declined with contact depth rather than being uniform (Figure S1). This null is conservative, however, because deep and cortical activity could be coupled; the cases that did exceed it therefore provide strong evidence of genuine deep sensitivity.

### High-frequency oscillations

Across our analyses, the correspondence between MEG and SEEG was weakest in the gamma range, while the low-frequency bands (delta, theta, alpha) exhibited the highest canonical correlation at depth. This is consistent with a signal-to-noise account: oscillatory power falls off steeply with frequency, so low-frequency activity sits well above the MEG noise floor, whereas gamma is intrinsically low in amplitude. Accordingly, the MEG-VE spectra were a flattened version of the SEEG spectra (Figure 2), the same pattern produced by adding white noise to the SEEG (Figure S2), suggesting that the MEG-VE behaves like a noisier measurement of the same underlying activity rather than capturing something qualitatively different.

The gamma-band activity recovered by previous simultaneous recordings was, however, task-evoked, and therefore transiently high in amplitude and synchronised across an extended cortical patch^19,25^. Ongoing gamma at rest is a harder target on both counts, and so a resting paradigm tests MEG under less favourable conditions than those earlier demonstrations. The weak correspondence we observe should not, therefore, be interpreted as a fundamental inability of MEG to capture gamma-band activity. Moreover, the limitations in detecting deeper and higher frequency signals that we have observed should be eased by on-scalp OPM-MEG, which places sensors closer to the brain and yields the largest gains for the weak fields produced by deep sources^31–33^. Applying the analyses developed here to OPM-MEG, ideally under a task designed to drive gamma-band activity, is a promising route to extending validated, non-invasive measurement to higher frequencies and to a wider range of deep structures^8,34^.

### Clinical and translational relevance

Previous studies suggest MEG and intracranial recordings agree on the localisation of epileptic activity, both at the seizure-onset zone^20,35^ and within the hippocampus^21^; our results extend this agreement from epileptiform discharges to spontaneous oscillatory dynamics. The whole-brain coverage of MEG complements the sparse, clinically-determined sampling of SEEG^36^, and non-invasive sensitivity to deep structures could help to target, or to reduce, invasive electrode placement^37^. Such applications are likely to become more practical with OPM-MEG, which is wearable and motion-tolerant and therefore well suited to paediatric and bedside use; on-scalp OPM-MEG has already been shown to evaluate epilepsy in school-aged children comparably to conventional MEG^6,38^.

More broadly, because MEG is non-invasive it can study deep oscillatory dynamics in contexts that intracranial recordings cannot reach: healthy participants, across the lifespan, and longitudinally. Non-invasive measurement of hippocampal theta, for example, would open the neural basis of memory to study in healthy people, building on prior MEG recordings of hippocampal theta during spatial learning^39^. Equally, validated measures of deep oscillatory activity could contribute to the early detection and monitoring of disorders such as Alzheimer’s disease, for which MEG has been proposed as a non-invasive biomarker^40^.

In addition, it is important to note that our recordings were made under atypical conditions for MEG: after electrode implantation, such that head dressing holds neural sources further from the sensors. In contrast, non-clinical and healthy volunteers can be positioned closer to the sensors and carry no such hardware. The correspondence we report is therefore likely to be a lower bound on what MEG can recover under typical recording conditions. It should also be noted that these are epileptic brains, and pathological activity may not be representative of the healthy populations to which we ultimately wish to generalise.

## Conclusions

Using a rare dataset of simultaneous MEG and SEEG recordings, we have established a direct, whole-brain benchmark for what source-reconstructed MEG can and cannot recover from spontaneous neural activity. Validated against an intracranial SEEG ground truth, MEG faithfully captured oscillatory power and transient bursts across the brain, including hippocampal theta, with the correspondence strongest for superficial regions and lower-frequency activity (delta, theta, and alpha) and weakest for gamma. These findings qualify the long-held view that MEG does not capture activity from deep structures, showing instead that its sensitivity declines with depth and frequency but remains physiologically meaningful. This work provides empirical validation for non-invasive markers that are widely used yet rarely tested against concurrent intracranial recordings, and a foundation for interpreting the rapidly growing body of MEG studies.

## Methods

### Data

#### Participants and acquisition

We analysed a simultaneous MEG-SEEG dataset acquired at rest from 14 participants (18-47 years; 7 female) with drug-resistant focal epilepsy, predominantly of temporal lobe origin, who were undergoing pre-surgical evaluation. The data were recorded at the Department of Neurosurgery, Affiliated Ruijin Hospital, Shanghai Jiao Tong University School of Medicine. See Zhang et al.^28^ for details regarding the acquisition setup. The study was approved by the local ethics committee of Ruijin Hospital and was conducted in accordance with the Declaration of Helsinki, and all participants gave written informed consent.

Intracranial depth (SEEG) electrodes were implanted under general anaesthesia for preresection seizure localisation, with the number and location of the electrodes determined solely by clinical indications. Each electrode carried 8 or 16 contacts (contact length 2mm, inter-contact spacing 1.5mm, contact diameter 0.8mm), so the number of contacts in each region varied across participants (Figure 1C). MEG was recorded simultaneously with the SEEG using a 306-channel Elekta Neuromag VectorView system in a lightly shielded room with active shielding. Both modalities were sampled at 1000Hz, and participants were instructed to rest with their eyes closed. The participant demographics are shown in Figure 1B and the SEEG contact counts and locations in Figure 1C.

The usable simultaneous-SEEG sample comprised 16 resting-state sessions from 14 patients, with some patients contributing more than one session. All analyses were computed per session.

#### Electrode localisation and regions of interest

The electrode contacts were localised using Lead-DBS; full details of the localisation procedure are given in Zhang et al.^28^. Each contact was assigned an Automated Anatomical Labelling (AAL3) anatomical label^41^ and a Montreal Neurological Institute (MNI) coordinate. For the region-level analyses, the AAL3 atlas was collapsed into a coarse 24-region scheme (10 bilateral cortical and subcortical groups, plus midline anterior cingulate, posterior cingulate, thalamus, and vermis).^1^

### Preprocessing

All preprocessing used osl-dynamics^42^ with MNE-Python^43^.

#### SEEG

After picking the SEEG channels, bad-channel detection was performed on the wide-band monopolar signal and bad contacts were dropped before referencing. The data were notchfiltered at 50 and 100Hz, band-pass filtered to 2 to 95Hz, and resampled to 250Hz. A bipolar montage was formed from adjacent same-shaft contacts less than 6mm apart; the anode was taken as the lower-numbered (more medial) contact, the bipolar location as the contact midpoint, and the region from the anode. Bad segments were detected on the bipolar signal using a generalized extreme Studentized deviate (ESD) procedure^44^, with four passes over the standard deviation, its temporal derivative, the kurtosis, and the derivative of the kurtosis (significance level 0.1, maximum fraction 0.3, 0.25s window), and low-variance bipolars (below the median minus three times the median absolute deviation of the log-variance) were dropped. Interictal epileptiform discharges (IEDs) were identified by visual inspection of the SEEG by a trained clinician. Marks were pooled across contacts, and a window of ±0.5s around each marked event was annotated as bad, so that all analyses exclude the same IED periods in both the SEEG and the simultaneously recorded MEG.

#### MEG

External magnetic interference was first suppressed using temporal signal-space separation (tSSS)^45^. The MEG data were then notch-filtered at 50 and 100Hz and band-pass filtered to 2 to 95Hz then resampled to 250Hz; bad segments and bad channels were detected using a generalized ESD procedure. Head surfaces were extracted from each participant’s T1-weighted MRI and the MEG was coregistered to the MRI using the Polhemus fiducials and headshape (RHINO;^46^).

### Source Reconstruction

#### Virtual electrodes

For each SEEG bipolar, we reconstructed a single VE using a unit-noise-gain LCMV beamformer^11^. The VE was placed at the bipolar midpoint and oriented along the bipolar (anode to cathode) axis. Source reconstruction used an MNE-Python volume source space and a singleshell boundary element model (conductivity 0.3) using the magnetometers and gradiometers. The data covariance was computed using the preprocessed data (excluding bad and IED segments; regularisation 0.01; rank 64) with an ad-hoc diagonal noise covariance using the average variance of each sensor type. For the per-band analyses, the data were band-pass filtered before beamforming to give one VE per session and band. Throughout, six frequency bands were used: delta (*δ*; 2 to 4Hz), theta (*θ*; 4 to 8Hz), alpha (*α*; 8 to 13Hz), beta (*β*; 13 to 30Hz), low-gamma (low-*γ*; 30 to 45Hz), and high-gamma (high-*γ*; 55 to 95Hz).

#### Random-location null virtual electrodes

To provide a spatially-mismatched control, we also reconstructed a set of random-location VEs. For each session, *K* = 100 points were sampled uniformly within the inner-skull surface, each with a random unit orientation, and a per-band LCMV beamformer was applied at each point. These VEs were used for the spatial-specificity null distributions described in the Statistical Analysis.

For the hippocampal analyses we additionally reconstructed a depth-matched control, which holds source depth fixed while varying location: 100 locations were sampled per session inside the inner-skull surface at the same distance-to-scalp shell as that session’s hippocampal contacts (within 5mm), at least 20mm from any hippocampal contact and outside the AAL hippocampus, each with a random unit orientation. It used the same LCMV settings as the random-location VEs, and was used for the spatial-specificity tests of the hippocampal power correlation (Figure S5) and amplitude modulation (Figure 4B).

### Cross-Modality Analyses

#### Canonical correlation analysis (CCA)

For each session, region, and frequency band, we computed the leading canonical correlation between the z-scored band-passed SEEG bipolars and the MEG. At the sensor level (Figure 1D), the MEG was the whole-head sensor array. At the source level (Figure S1A), the matched MEG-VE set was used. Significance was assessed by block-shuffling permutation (see Statistical Analysis), and we report the bias-corrected effect *r*_eff_ = *r*_obs_ − the null mean, together with *p*_perm_. The depth of each contact was defined as the Euclidean distance from its midpoint to the nearest point on the fsaverage scalp surface, and the region depth as the mean over its contacts; the headline statistic is the Spearman correlation between *r*_eff_ and depth across regions for each band. Only regions that had at least 3 sessions with at least 10 SEEG contacts in total were included.

#### Oscillatory power

Before spectral estimation, each SEEG contact and MEG-VE channel was z-scored within session, so that estimates are comparable across participants. The spectra are therefore in normalised units (fractional variance per Hz), and only their shape is interpretable, not their absolute values. PSDs were estimated using Welch’s method (4s segments, Hann window, 50% overlap), and band power was taken as the integral of the PSD over each band. Each region was summarised as the log10 of the median power across its contacts, and PSDs were averaged across sessions as the (geometric) mean log10 power with the standard error of the mean.

Cross-modal agreement (Figure 2) was quantified, for each region and band, as the Pearson correlation across sessions (with the region filter as above); the per-band value annotated in the figure pools all (session, region) points. These points are treated as independent observations, which they are not strictly: a session contributes up to ten regions, and two patients contributed two sessions each. The pooled correlation is therefore reported as a descriptive summary of the relationship rather than as a formal test, and the corresponding inferential claim rests on the random-VE null (Figure S3), which is computed through the identical pooling structure and so is unaffected by this dependence.

#### Oscillatory bursts

Oscillatory bursts are transient, high-amplitude events^22,23,47^. They were detected per contact and band by band-pass filtering (4th-order Butterworth, forwards-backwards) and taking the Hilbert amplitude envelope, then thresholding at the 80th percentile of that contact’s within-session envelope; supra-threshold runs separated by at most one cycle (at the band centre) were merged, runs shorter than three cycles were discarded. Contacts whose mean band envelope was below half the session median were excluded.

To quantify the SEEG amplitude modulation accompanying MEG bursts (Figure 3), the MEG-VE burst train was used to split the simultaneous SEEG envelope into in-burst and outof-burst periods, and we computed the modulation index:

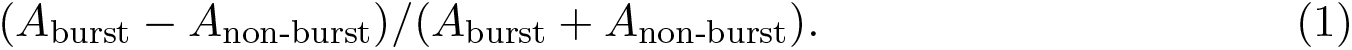

The existence of modulation was assessed by block-shuffling permutation (see Statistical Analysis), giving a per-contact *z*-score and one-sided *p*-value. Values were aggregated to one point per session and region (at least 3 contacts, with the region filter as above), and the marker fill indicates a Stouffer-combined *p* < 0.05.

The same procedure was applied to the hippocampus (Figure 4B). Temporal coupling was assessed, per band, as a one-sided Wilcoxon signed-rank test of the per-session modulation against zero. Spatial specificity was assessed by gating the same SEEG envelope with bursts detected on the matched-depth control VEs, and comparing the true-location modulation to the matched-depth modulation across sessions (paired one-sided Wilcoxon).

### Statistical Analysis

The significance of the cross-modal correspondence (the canonical correlations and the amplitude modulation, both reported as *p*_perm_) was assessed using a block-shuffling permutation test, in which the signal was divided into 1s blocks that were randomly shuffled to generate a null distribution, breaking the temporal alignment between MEG and the simultaneously recorded SEEG while preserving the within-block temporal structure. For the canonical correlations, the MEG was shuffled (*N* = 100 permutations); for the amplitude modulation, the SEEG envelope was shuffled while the MEG burst train was held fixed (*N* = 500).

Spatial specificity was tested using the random-location VEs: for the power correlation (Figure S3) the MEG band powers were replaced by random-VE draws (*N* = 200) to give a per-band Monte-Carlo *p*-value (corrected using the Benjamini-Hochberg false discovery rate, BH-FDR); and for the amplitude modulation (Figure S4) the burst train was instead taken from random VEs in a different region, giving a per-contact *z*-score that was Stouffer-combined. For the hippocampal power correlation (Figure S5) the same Monte-Carlo procedure was applied using the matched-depth VEs in place of the uniform random-location VEs, replacing the MEG-VE band powers by *N* = 200 depth-matched draws and recomputing the cross-session correlation. Benjamini-Hochberg FDR^48^ was applied for Figure 2B, Figure S3, and Figure S5.

## Acknowledgements

We thank Neil Burgess and Eleonora Marcantoni for their useful comments on the manuscript.

G.B. was supported by the Discovery Research Platform for Naturalistic Neuroimaging funded by Wellcome (226793/Z/22/Z). V.L. was supported by a Royal Society International Exchanges grant (IEC\NSFC\211206). M.W. was supported by the Wellcome Trust (106183/Z/14/Z, 215573/Z/19/Z), the New Therapeutics in Alzheimer’s Disease (NTAD) study supported by the Medical Research Council, the Dementia Platform UK (RG94383/RG89702) and the NIHR Oxford Health Biomedical Research Centre (NIHR203316). C.C. was supported by the National Natural Science Foundation of China (NSFC) (Grant No. 82071547) and University College London-Shanghai Jiao Tong University Strategic Partner Funds (2023). D.B. was supported by a UKRI Frontier Research Grant (EP/X023060/1). U.V. was supported by the Wellcome Trust (307306/Z/23/Z) and the Engineering and Physical Sciences Research Council (EP/Z533191/1).

## Author contributions

C.G. contributed to conceptualization, methodology, validation, formal analysis, investigation, visualization, writing (original draft) and writing (review and editing). G.O. contributed to data curation and writing (review and editing). G.B. contributed to methodology. V.L. contributed to writing (review and editing). M.W. contributed to methodology and writing (review and editing). S.Z., W.L. and B.S. contributed to investigation and resources. C.C. contributed to resources and project administration. D.B. contributed to conceptualization, methodology, supervision, writing (original draft) and writing (review and editing). U.V. contributed to writing (review and editing) and supervision.

## Footnotes

1 Details of this 24-region AAL3 parcellation are available at https://osl-dynamics.readthedocs.io/en/latest/parcellations/aal24.html.

