## Supplementary information for "Intracranial Validation of Magnetoencephalography Across Oscillatory Frequency and Depth"

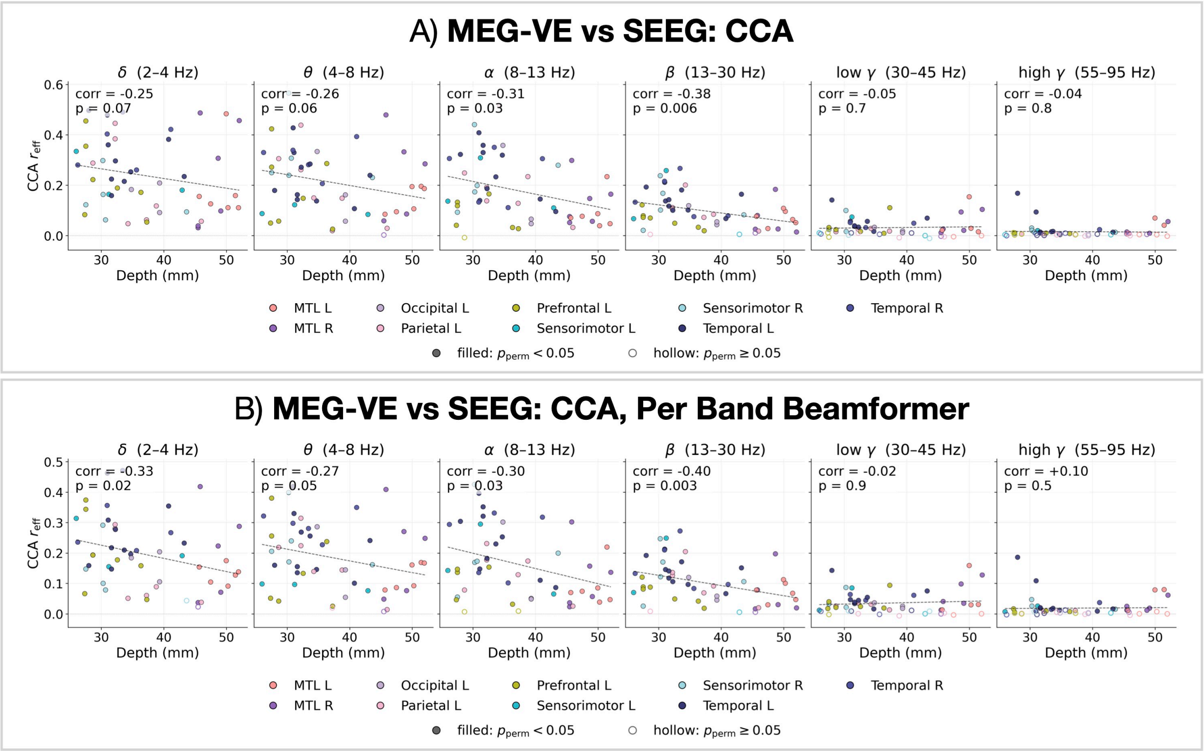


Figure S1: **MEG-VE vs SEEG: canonical correlation analysis**. A) Canonical correlation between MEG virtual electrodes (MEG-VE) and SEEG, plotted against mean contact depth for six frequency bands (the VE counterpart of Figure 1D, which uses MEG sensors). Each point is a region in one session; filled points denote a significant canonical correlation (block-permutation test, $p_{\text{perm}} < 0.05$) and the annotated corr/$p$ give the relationship between CCA and depth. B) The same analysis using a separate beamformer optimised for each frequency band, which slightly improves the correspondence. As for the sensor-level analysis, correlations are strongest for superficial contacts and low frequencies.

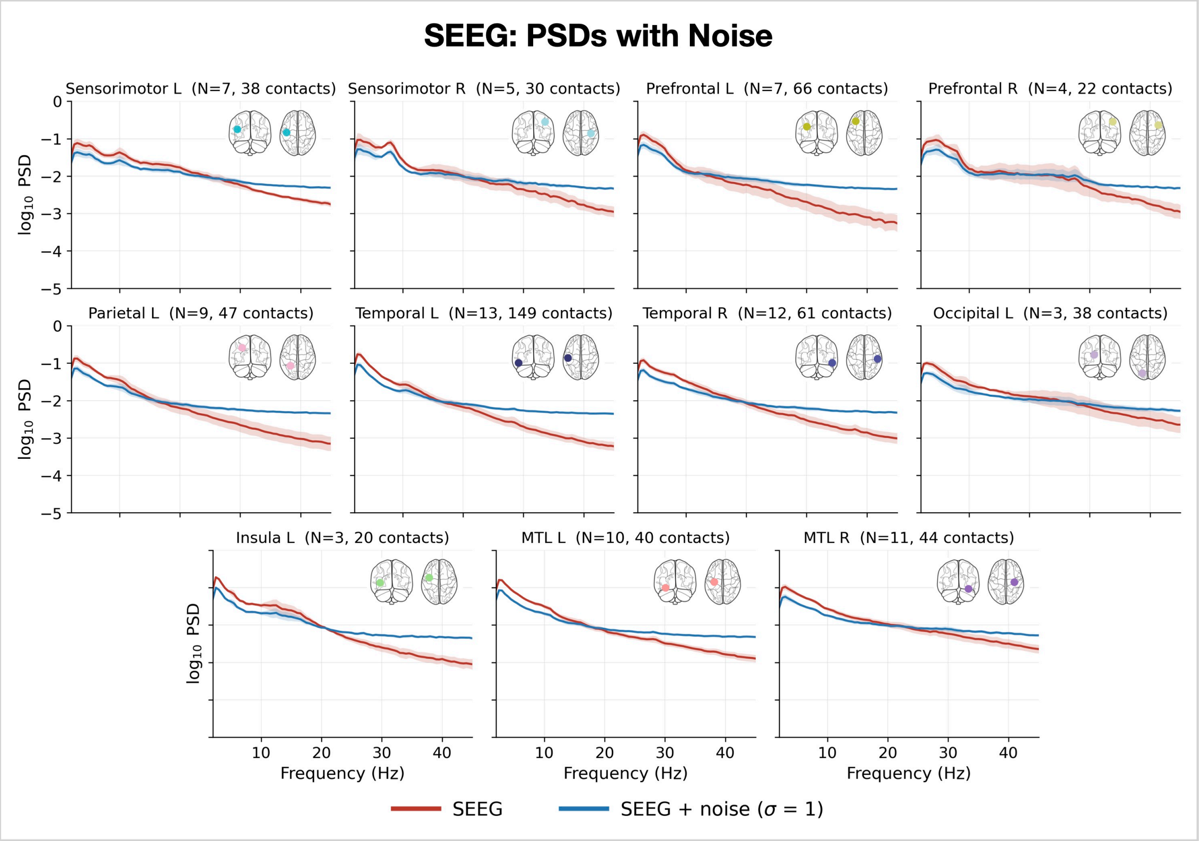


Figure S2: **SEEG PSDs with added noise**. Power spectral density (normalised units, fractional variance per Hz) for each region computed from the SEEG (red) and from the SEEG after adding white noise ($\sigma = 1$; blue), averaged across sessions and contacts (line, mean; shaded, variability). Adding noise flattens the high-frequency SEEG spectrum and reproduces the spectral shape of the MEG-VE (cf. Figure 2A), consistent with MEG behaving like a noisier measurement of the same underlying activity. Because each channel is z-scored before spectral estimation (spectra in fractional variance per Hz), the added noise raises the total variance and thereby redistributes normalised power towards higher frequencies; the absolute low-frequency power is unchanged, but its share of the now-larger total is reduced, producing the apparent flattening.

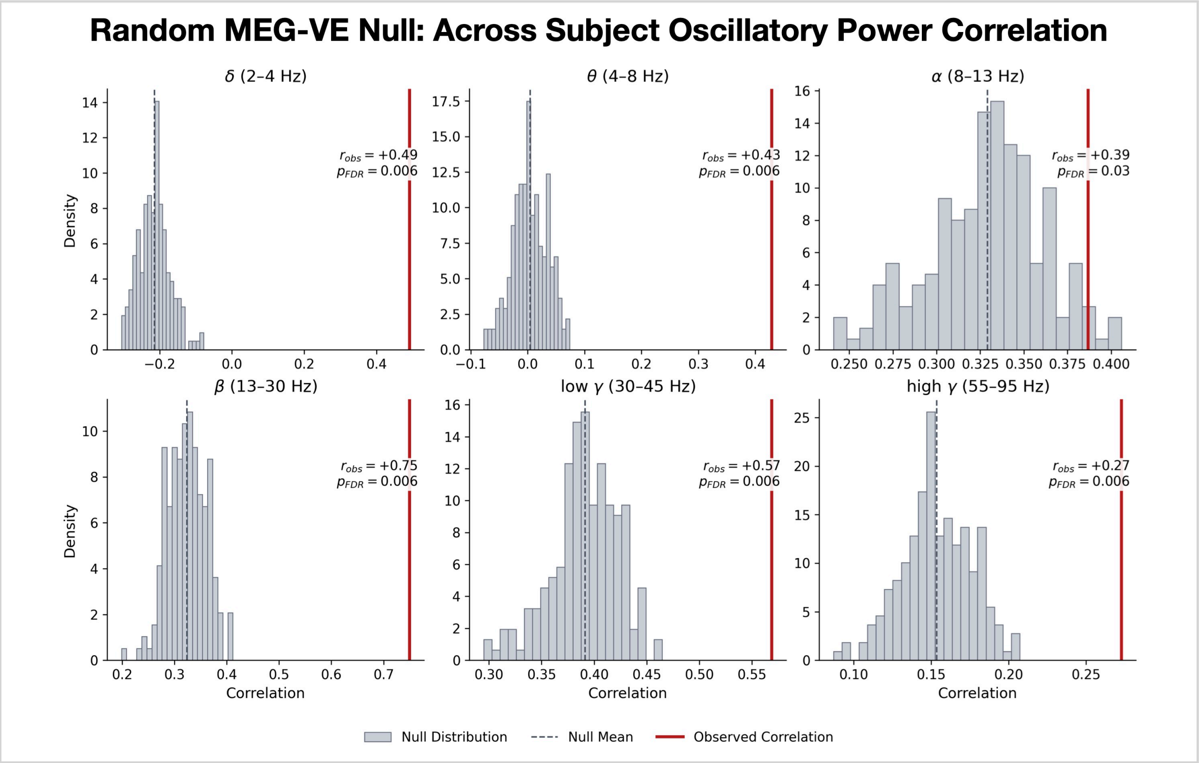


Figure S3: **Spatial specificity of the oscillatory power correlation**. Null distributions (grey histograms) for the across-session correlation between MEG-VE and SEEG oscillatory power, obtained by reconstructing the MEG-VEs at random (spatially mismatched) locations, for each frequency band. The observed correlation (red line; $r_{\mathrm{obs}}$) exceeds the null in every band (dashed line, null mean; $p_{\mathrm{FDR}}$, FDR-corrected), indicating that the MEG-VE-SEEG power correspondence is spatially specific.

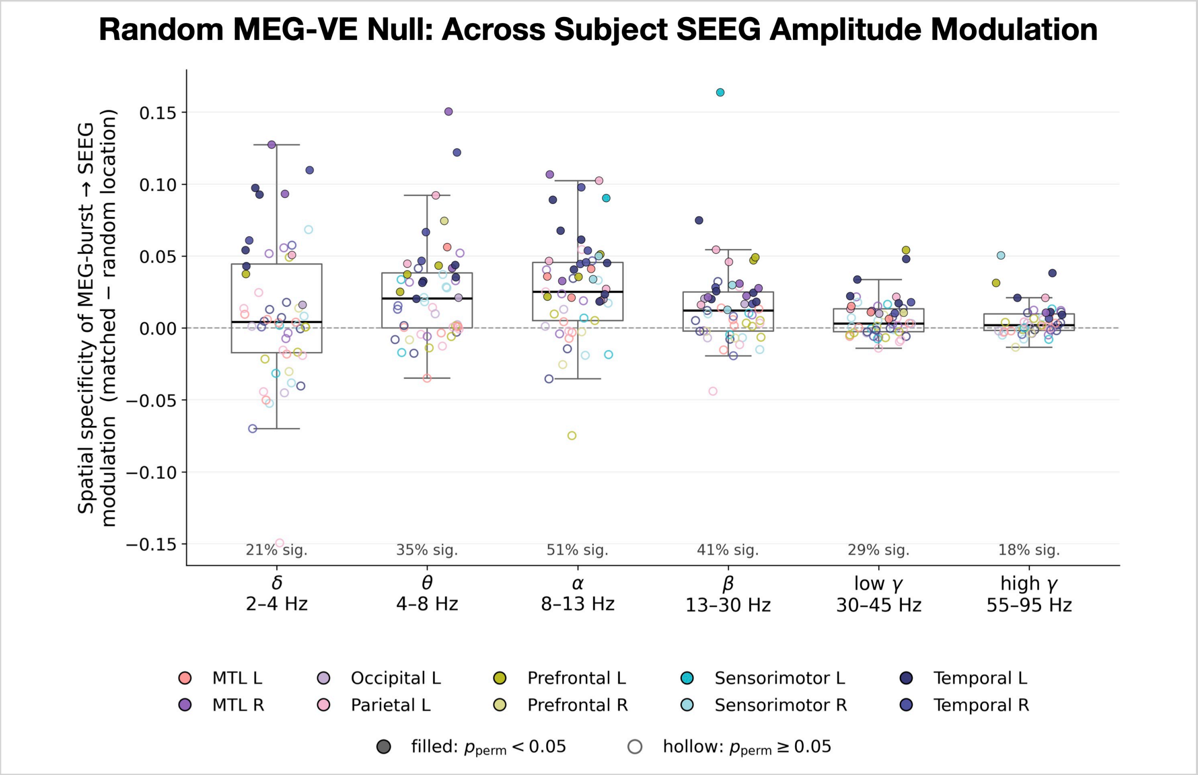


Figure S4: **Spatial specificity of the SEEG amplitude modulation**. Distribution across sessions of the difference in SEEG amplitude modulation index between the true MEG-VE and a null in which the MEG-VEs are reconstructed at random (spatially mismatched) locations ($\Delta$ mod. index), for each frequency band. Positive values indicate stronger modulation at the true location; the percentage of sessions with a significant difference is annotated and peaks in $\theta/\alpha$. Points are coloured by region and filled points are individually significant (block-permutation test, $p_{\text{perm}} < 0.05$).

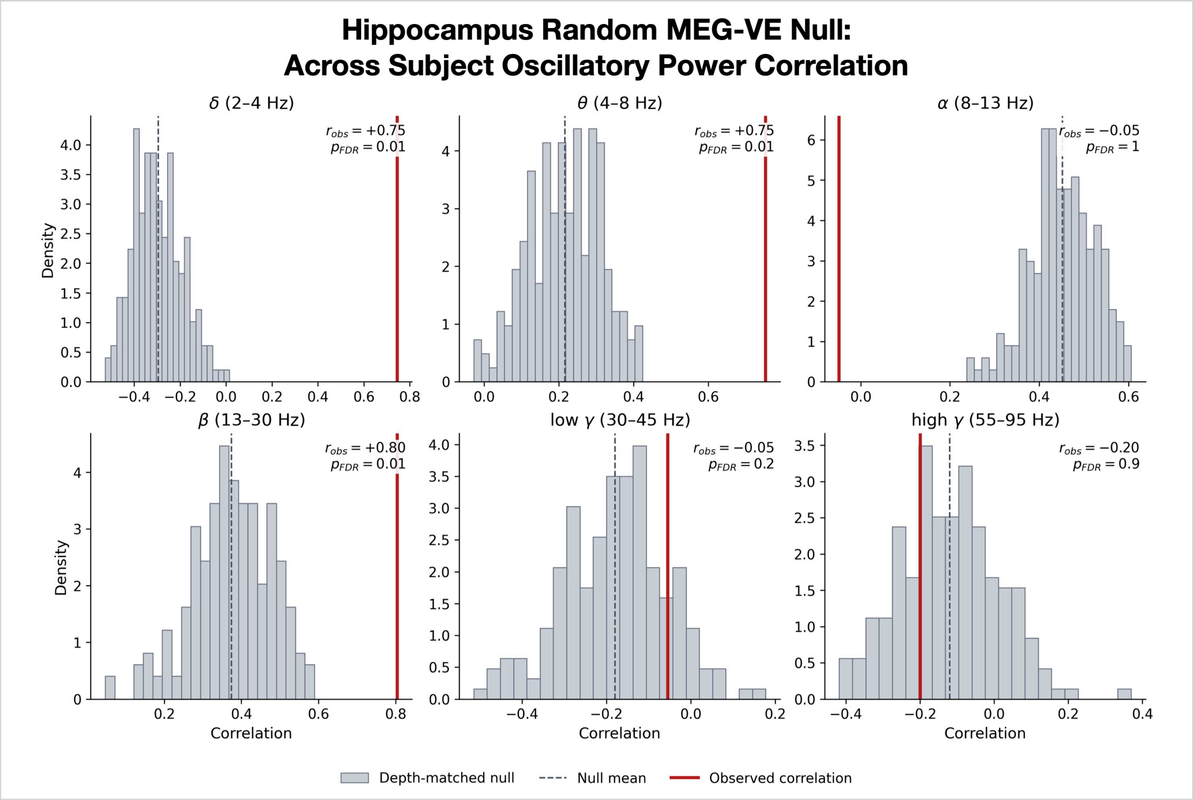


Figure S5: **Spatial specificity of the hippocampal oscillatory power correlation**. Null distributions (grey histograms) for the across-session correlation between MEG-VE and SEEG hippocampal oscillatory power, obtained by reconstructing the MEG-VEs at depth-matched locations (sampled at the same distance-to-scalp shell as the hippocampal contacts, at least 20 mm away and outside the hippocampus) for each frequency band.

### Functional connectivity

Our participants were an epilepsy population with clinically-placed electrodes, which introduces the possibility of pathology and results in spatially biased and sparse coverage. This sparse, clinically-determined sampling prevented an accurate estimation of functional connectivity, which we therefore did not analyse in detail; for completeness, we report an amplitude envelope correlation analysis in Figure S6, which shows cross-modal correspondence for those edges that could be estimated.

**Amplitude envelope correlation (AEC)** was computed per session at the contact level and then aggregated to a region-by-region connectome (the mean over inter-region contact pairs, with the within-region diagonal dropped and an edge retained only if it was present in at least 3 sessions). For the SEEG, the data were band-pass filtered (4th-order Butterworth), Hilbert-enveloped, z-scored, and correlated (Pearson) over 30 s blocks; no orthogonalisation was applied, as the bipolar referencing already removes most volume conduction. For the MEG-VEs, the per-band VEs were pairwise-orthogonalised[49] to remove zero-lag source leakage before the same 30 s block-averaged correlation, which was averaged over the two orthogonalisation directions. Cross-modal agreement (Figure S6) was assessed by aggregating both modalities to region matrices (diagonal dropped), taking the upper-triangular edges as paired (SEEG, MEG) points, and computing the Pearson correlation for each band.

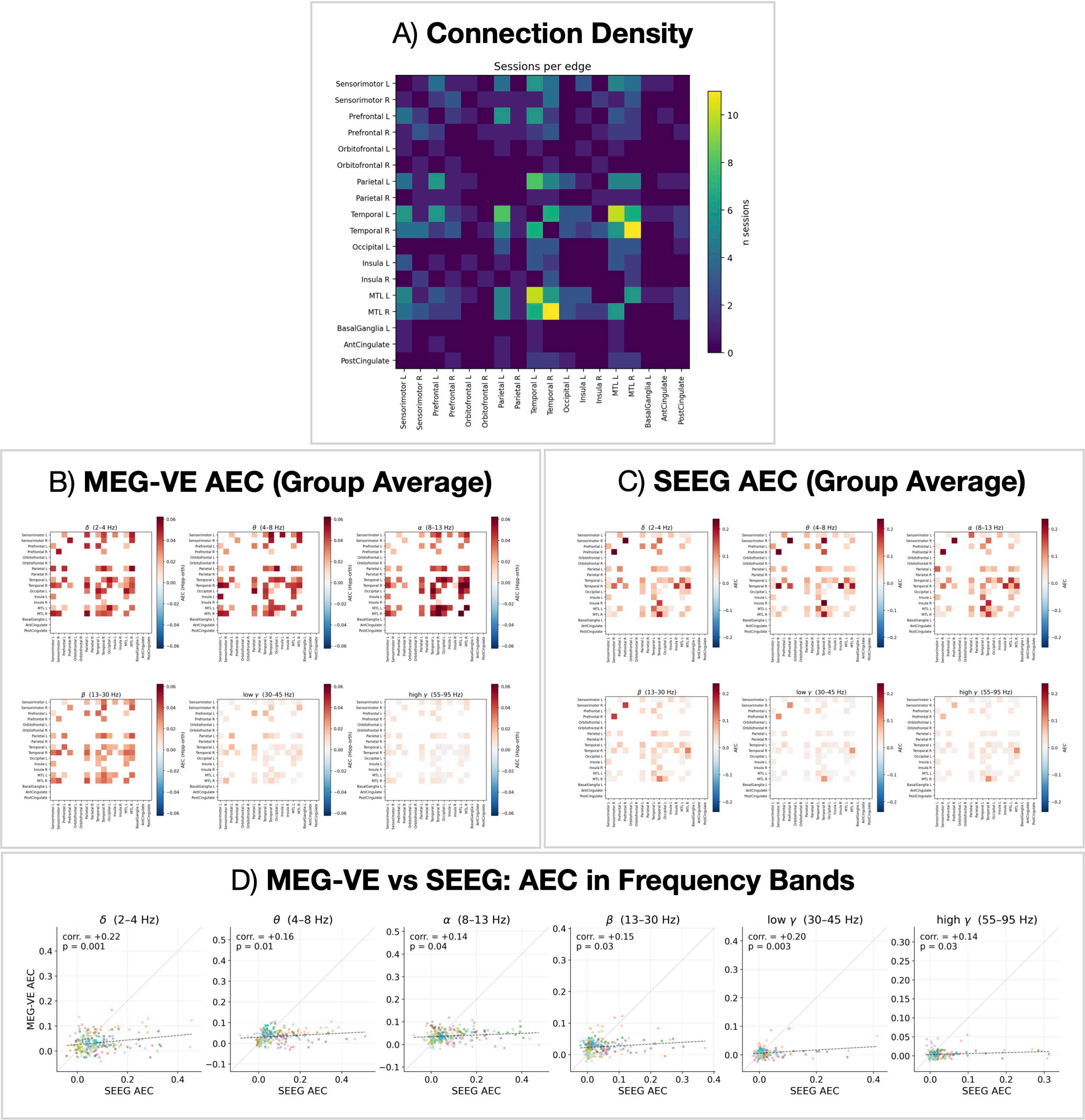


Figure S6: **AEC analysis**. A) Connection density: the number of sessions in which each region-pair edge could be estimated (i.e. both regions had SEEG contacts), which limits the network coverage. B) Group-average MEG-VE AEC and C) group-average SEEG AEC, shown per frequency band. D) Across-session correlation between MEG-VE and SEEG AEC for each frequency band (each point an edge; dashed line, line of best fit).
